# Trophic resources of the edaphic microarthropods: a worldwide review of the empirical evidence

**DOI:** 10.1101/2021.02.06.430061

**Authors:** Víctor Nicolás Velazco, Leonardo Ariel Saravia, Carlos Eduardo Coviella, Liliana Beatriz Falco

**Affiliations:** Departamento de Ciencias Básicas, Universidad Nacional de Luján (Argentina), Ushuaia, Argentina; Instituto de Ecología y Desarrollo Sustentable (UNLu – CONICET), Ushuaia, Argentina; Instituto de Ciencias, Universidad Nacional de General Sarmiento (Argentina), Ushuaia, Argentina; Centro Austral de Investigaciones Científicas (CADIC-CONICET), Ushuaia, Argentina

**Keywords:** food web, trophic ecology, soil mesofauna, Acari, Collembola

## Abstract

Ecosystem sustainable use requires reliable information about its biotic and abiotic structure and functioning. Accurate knowledge of trophic relations is central for the understanding of ecosystem dynamics, which in turn, is essential for food web stability analyzes and the development of sustainable practices. There is a rapid growth in the knowledge on how belowground biodiversity regulates the structure and functioning of terrestrial ecosystems. Although, the available information about trophic relationships is hard to find and fragmented. This gathering the information available worldwide about the food resources of soil mesofauna. From the 3105 hits of the initial search on food resources of soil microarthropods, only a total of 196 published works related particular species, genera, and families to particular trophic resources, the majority of them dealing with soils of the Palearctic region. From the 196 publications we extracted 3009 records relating specific taxonomic groups to their trophic resources, 20% mention saprophytic fungi as a food resource, 16% cite microfauna, 11% mention bacteria, 10% litter and 5% cite Mycorrhizal fungi. The available information was highly skewed, the 73.71% comes from Acari, and within these, 50.62% correspond just to Sarcoptiformes. For Collembola, the literature is scarce, the majority coming from Arthropleona. This review highlights the general lack of information relating species, genera, and families of the soil mesofauna to specific trophic resources. It also highlights that available research mostly comes from European sites, with the use of trophic resources by the mesofauna of the majority of the soils in other parts of the world still largely unknown.

## Introduction

Only in the last few decades, sustainable use of the soil has come sharply into focus. The report on Global Biodiversity and Ecosystem Services (IPBES, 2019), and the recent State of Knowledge of Soil Biodiversity (FAO, 2020) clearly state the importance of the soil ecosystem and the central role that soil biodiversity plays on ecosystem services. However, critical information is lacking on edaphic biodiversity and interaction webs for most soils.

The view about the edaphic biota has changed in recent years. Much of the work on taxonomic descriptions, richness, biodiversity estimation, and microenvironments (Anderson, 1978; Petersen and Luxton, 1982), highlights the fragility and need for the conservation of soils, and soil biodiversity (FAO, 2020; Walker, 1992). The focus has gradually shifted towards the study of the functioning of the soil ecosystem, in which the interactions between the biota and the edaphic environment hierarchically affect its structure and functioning, regulating the functioning of the edaphic ecosystem. (Barrios, 2007; Behan-Pelletier and Newton, 1999; Brussaard, 1997; Lavelle et al., 2006). The relationships of species richness with ecosystem dynamics allow establishing causal relationships between the characteristics of the organisms present and the processes and services of the ecosystem (Hooper et al., 2005; Martín-López et al., 2007) in natural as well as in managed environments. (Wall et al., 2012).

The soil mesofauna is mainly constituted by microarthropods belonging to the Subclasses Acari and Collembola (following Kranz and Walter, 2009; Hopkin 2007) that inhabit the upper soil horizons (Martinez and Narciso, 2009), where complex communities are developed and maintained. These communities are influenced by a high microhabitat heterogeneity and the number of food resources available (Anderson, 1977, 1978; Lavelle and Spain, 1994), which provide a wide and varied set of ecological niches (Nielsen et al., 2010; Wallwork, 1958). The mesofauna, through its trophic relationships, contributes to the edaphic functioning through the fragmentation of organic matter, the nutrient cycle dynamics, the transport of microflora propagules, and the regulation of microflora and microfauna populations that affect primary production. (Brussaard, 1997; Cragg and Bardgett, 2001; Lavelle, 1996; Wall et al., 2012; FAO, 2020).

Studies on the flow of energy and matter in the soil ecosystem, group edaphic organisms into functional guilds to understand the relationships between soil biota structure and ecosystem functioning. However, this approach leaves aside the understanding of how changes in the community influence the ecosystem’s functioning (Thompson et al., 2012; Wall and Moore, 1999). Thompson et. al. (2012) suggests addressing this problem from the point of view of the trophic networks, which could link the flow of matter and energy and the diversity of a community. Thus, by studying the structure of the trophic networks, we can better understand the role of biodiversity in the functioning of the ecosystem.

Knowing the resources that microarthropod groups use for food is difficult due to their size and cryptic environment. The recognition of the food resources that are part of the diet of soil microarthropods must be based on empirical evidence, which constitutes the first step to establish the interactions between trophic species and the trophic resources that describe the food webs. (Briand and Cohen, 1984; van Straalen, 1998; Walter et al., 1991).

The information about trophic relationships of the different taxonomic groups is still quite scarce. These data are necessary to build and analyze webs of interactions that, in turn, could be used to assess the stability and conservation status of these ecosystems. (Barrios, 2007; Briones, 2014; Thakur et al., 2020).

This review gathers the information currently available regarding the use of trophic resources by the edaphic microarthropods. This information, together with other characteristic features of the soil microarthropods, will allow building a food web interactions network, that could result in a better understanding of the structure and functioning of the edaphic biota (FAO, 2020). Trophic networks will, in turn, allow for comparing the state of different soils, or the same soils under different intensities of anthropic impact (Thompson et al., 2012).

The empirical characterization of trophic interactions is challenging due to the spatial and temporal complexity of feeding patterns, and the limitations of the methods to identify and quantify the components of the diet (Nielsen et al., 2018; Pankhurst et al., 1997; Walter et al.,1991).

Thus, the objectives of this review are: 1-to gather all the information currently available regarding the trophic resources used by soil microarthropods, 2-to describe the known trophic relationships and potential diets of these soil microarthropods at the family taxonomic level or lower, and 3-to establish the current status of knowledge, skews in the available information.

## Materials and methods

### Empirical evidence

The empirical evidence provided through studies carried out under laboratory conditions (A) can be based on **observations** of the feeding behavior of the animals under study, on studying **food preferences**, or **tests** to study other interactions or biological phenomena related to diet, in general, the tests they are made using microcosm (Saur and Ponge 1988; Hubert, Žilová, and Pekár 2001; Schneider 2005; Schneider and Maraun 2009). The observation of the **intestinal content** (B) is based on considering that what is found in the tract is evidence of what is actually consumed (Jacot in 1936; Hartenstein 1962; Schneider et al. 2004; John A. Wallwork 1958a) this method requires the preparation of specimens for observation by microscopy techniques. The morphology and functioning of the **mouthparts** (C) are related to the manipulation, acquisition, and processing that microarthropods carry out on food and could be used to determine trophic guilds (Buryn and Brandl 1992; Kaneko 1988; Macnamara 1924; Poole 1958). This evidence also requires microscopy techniques to obtain information.

**Molecular** tools (D), such as barcoding, could be used to determine the presence of a species or a taxonomic group within the intestinal tract and it allows for establishing trophic relationships or other types of interactions. (Heidemann et al. 2014; King et al. 2008; Read et al. 2006). The study of **digestive enzymes** (E) in invertebrates can explain the digested food portions (Siepel and Ruiter-Dijkman 1993; Berg, Stoffer, and Van Den Heuvel 2004), those enzymes that hydrolyze structural polysaccharides are related to the diet (Nielsen 1962) and allow the differentiation of trophic guilds (Berg, Stoffer, and Van Den Heuvel 2004; Siepel and Ruiter-Dijkman 1993).

The use of the natural variation of **stable isotopes** (F) as empirical evidence of the use of trophic resources is based on the study of isotopic signatures; the isotopic signature of 15N informs about the trophic level of the invertebrate and that of 13C will indicate the proportion of trophic resources consumed, but the potential trophic resources need to be chosen previously to the isotopic analysis. (Tiunov 2007; Maraun et al. 2011; Potapov, Tiunov, and Scheu 2019; Potapov et al. 2019) also, the potential of this tool is such that inferences can even be made about the metabolic pathways of biomolecules (Pollierer et. Al. 2019, Pollierer & Scheu 2021; Chamberlain et. Al. 2006).

Finally, the study of the **lipid profile** (G) is based on using the fatty acids of the tissue of the resources as biological markers through the identification of fatty acids in the animal, since it or cannot synthesize them (absolute markers) or their synthesis supposes a high metabolic cost (relative markers) (Kühn, Schweitzer, and Ruess 2019; Ruess and Chamberlain 2010; Sechi et al. 2014; Paul M. Chamberlain and Black 2005; Pollierer, Scheu, and Haubert 2010).

General review publications are also included in this review, provided that they collect information on the use of trophic resources from publications not readily available.

### Bibliography search

A systematic search for empirical evidence that relates the use of at least one trophic resource by soil microarthropods is the strategy used to address this issue. For them, we relied on publications whose keywords relate to the taxonomic groups of interest, trophic relationships, and methods that provide evidence of consumption. From these keywords, we formulated search strings applied to scientific bibliography in both Scopus and Google Scholar database search engines.

#### We used the following search string in Scopus

“ALL((microarthropods OR springtails OR mites OR oribatida OR mesostigmata OR prostigmata OR astigmata) AND (trophic OR diet OR feeding) AND soil AND (“gut content” OR “stable isotope” OR “food preference” OR “fatty acid” OR lipids OR metabarcoding) AND (family OR genus OR species)) AND (LIMIT-TO (SUBJAREA, “AGRI”) OR LIMIT-TO (SUBJAREA, “ENVI”) OR LIMIT-TO (SUBJAREA, “MULT”) OR LIMIT-TO (SUBJAREA, “EART”)) AND (LIMIT-TO (EXACTKEYWORD, “Collembola”) OR LIMIT-TO (EXACTKEYWORD, “Acari”) OR LIMIT-TO (EXACTKEYWORD, “Soil Fauna”)) AND (LIMIT-TO (LANGUAGE, “English”) OR LIMIT-TO (LANGUAGE, “Spanish”))”, which returned 838 titles (September 3, 2021).

#### For searching in Google Scholar we used

“(microarthropods OR springtails OR mites) AND (“oribatida” OR “mesostigmata” OR “prostigmata” OR “astigmata”) AND (trophic OR diet OR feeding) AND soil AND (“gut content” OR “stable isotope” OR “food preference” OR “fatty acid” OR metabarcoding)”. This search returned 2170 titles (September 3, 2021).

#### We made additional searches from reviews and books resulting in 97 more titles

The first eligibility criterion to reduce the number of records was to select those papers whose titles or abstracts relate soil microarthropods to trophic resources (425 titles). From these, we selected those papers that effectively mention families, genera, or species of soil mites and springtails, relating them with at least one trophic resource. This resulted in 196 publications, which can be consulted in Electronic Supplementary Material I (ESM I). All the microarthropods recorded in the 196 publications cited can be found in ESM II, which is cross-referenced with ESM I. A flow diagram of the bibliography search and selection procedure is also provided as ESM III.

### Database construction

The empirical evidence provides information on the trophic relationships of these animals in different ways: some publications relate the taxa to food items and others group them into guilds or trophic categories. To deal with this heterogeneity, it was necessary to define the trophic resources (Table 1) that summarize their trophic and ecological characteristics in the edaphic ecosystem (Berg & McClaugherty 2008; Clark 1971; Warcup 1971; Ponge 1991; Persson et al. 1980; Krantz et al. 2009; Chernova et al. 2007; Rusek 1998; Schneider et. Al. 2005).

**Table 1:**
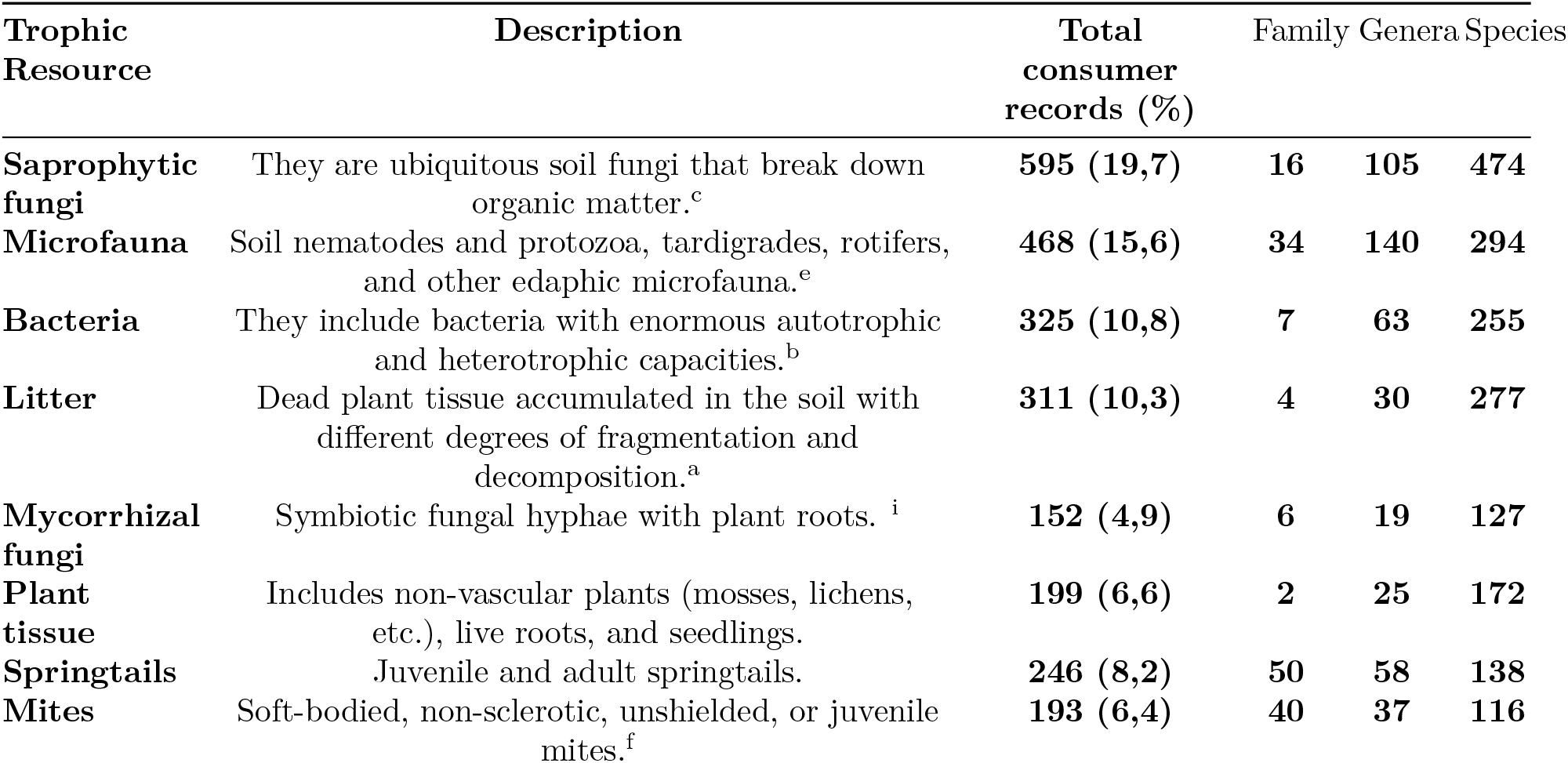
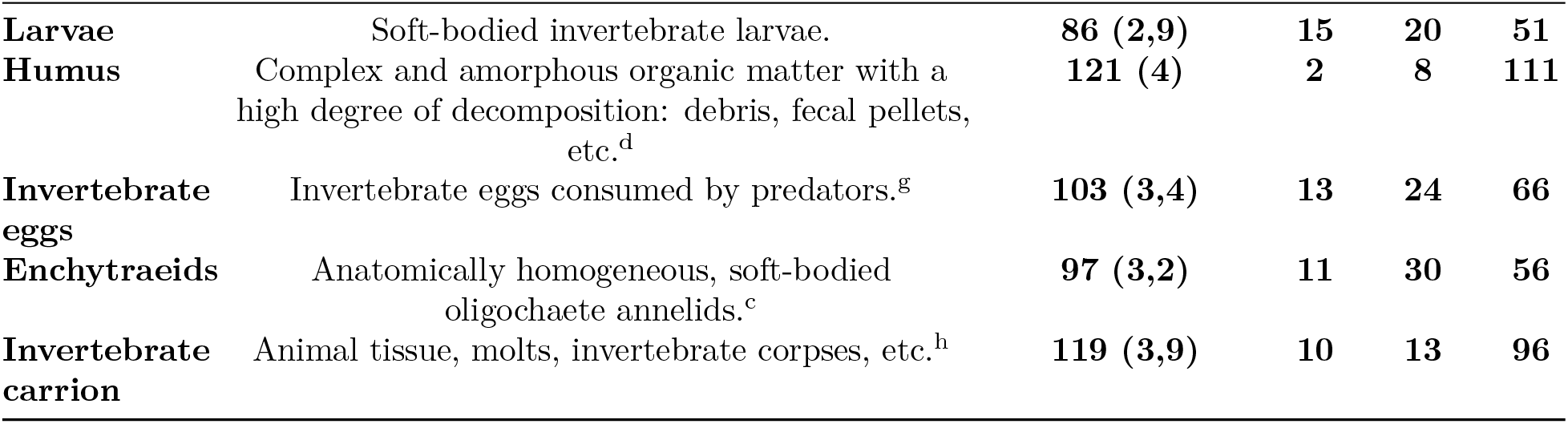
Basic description of trophic resources. Total records of consumers by trophic resource with absolute values and percentages; and the number of records by taxonomic resolution.^a^Berg & McClaugherty 2008 ^b^Clark 1971 ^c^Warcup 1971 ^d^Ponge 1991 ^e^Persson et al. 1980 ^f^Krantz & Walter 2009 ^g^Chernova et al. 2007 ^h^Rusek 1998 ^i^Schneider et al. 2005

We associate taxa and those resources, so that: a) if the publication indicated a food item it was assigned to a trophic resource in which it is defined, b) or if the taxa are grouped in some guild or trophic category, then each taxon was assigned the typical resources consumed by that category.

Based on this allocation strategy, we developed a database presented in Supplementary material II. The taxonomic information and the trophic resources were obtained from the different sections of the publications and their appendices (Thakur et al., 2020). In a complementary way, each taxon found has all the classification levels according to Krantz 2009 and Hopkin 2007, for mites and springtails respectively. The database is also available in the github repository https://github.com/EcoComplex/TrophicResources and Zenodo https://doi.org/10.5281/zenodo.6508661.

### Data analysis

The records were then analyzed by counting the associations between the taxa, the methods used and the trophic resources identified. We identified trophic resources, the number of records, and their relationships with the different taxonomic levels. Then, the breakdown of each taxon was carried out within the taxonomic categories and their relationship with trophic resources. We calculated the proportions of the different methods and their relationship with the taxonomic level and trophic resources.

Finally, we estimated the potential importance of resources on the diet for the main orders of mites (Arachnida) and springtails (Collembola) using the proportion of mentions between trophic resources and taxa included in such orders. For example, if a species has ten records on the consumption of the same trophic resource, the selection of this food item is likely a reflection of the use of the resource. Then if we gathered the information available from different species of the same genus, and the food items consumed by them, then we could assign it to the potential diet for the genus. Similarly, the resource use of the genera within a family can be thought of as reflecting the potential diet of the family.

That is, the diet of higher taxonomic hierarchies will be constituted by the sum of the resources used by the lower taxonomic levels.

The calculations, graphs, and tables were prepared using Microsoft Excel and R Statistical software version 4.1.2 (R Core Team 2021), the source code is available at Github https://github.com/EcoComplex/TrophicResources and Zenodo https://doi.org/10.5281/zenodo.6508661.

## Results

From the 3105 papers initially recovered with the searches, 196 papers were found to meet our criteria (see ESM III). A total of 133 articles from the 196 selected publications (ESM I) mention the countries in which the studies were carried out. Of these, 1 article belongs to the Ethiopian region, 3 to the Neotropical region, 3 to the Oriental region, 7 to the Australian region, 34 to the Nearctic region, 75 are all located in the Palearctic region, and the remaining 10 are from the Antarctic region. (Figure 1). This is important to highlight because the species strongly vary depending on the biogeographic region. There are 106 publications mentioning the environments in which the studies were conducted; from these 59 were temperate forests, 3 tropical forests, 21 grasslands, 23 agroecosystems, and 11 deserts.

**Figure 1:**
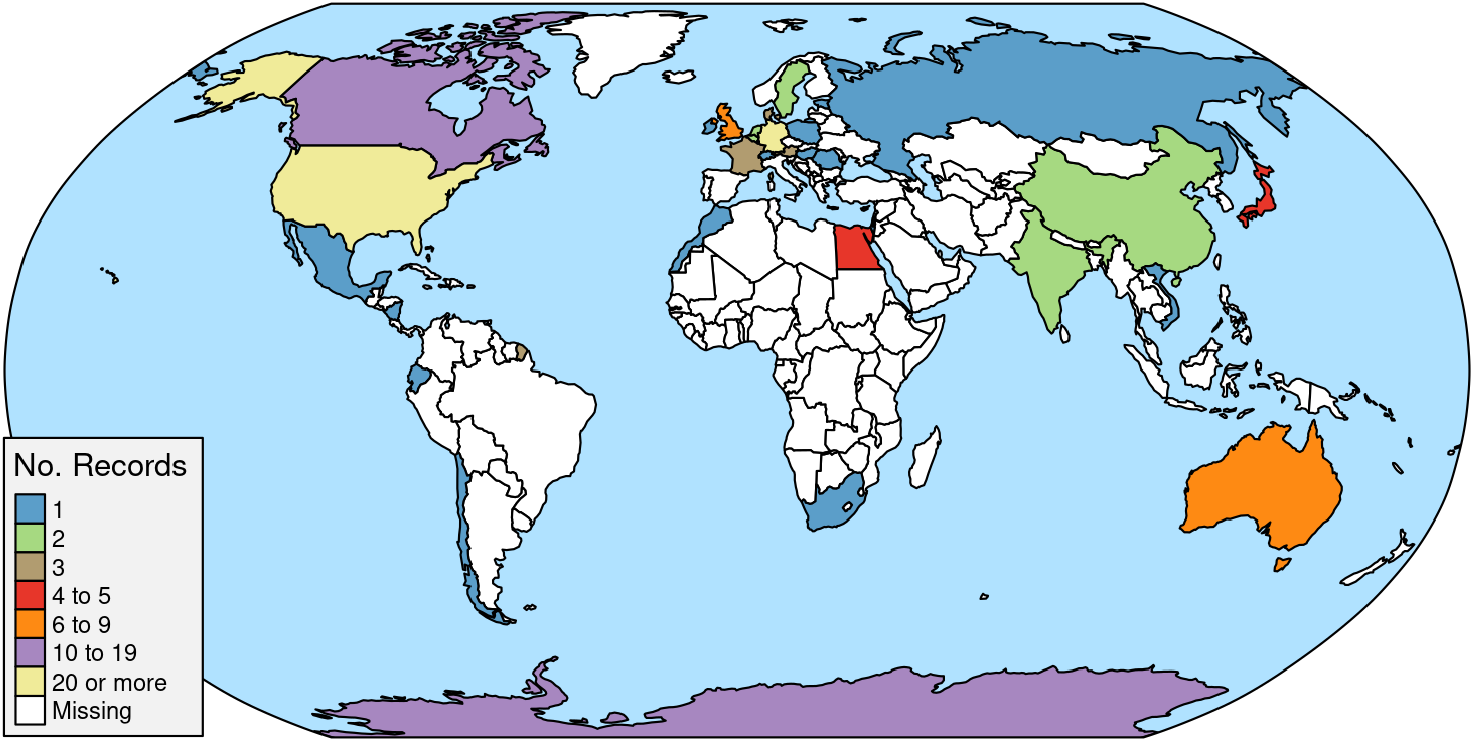
World map showing the distribution of records assigning trophic resources to soil mesofauna, largely unexplored outside Europe and the United States.

We obtained a total of 3015 records on trophic relationships (ESM II), with Acari contributing 2218 records (73.57%), and within this number, the majority (50.86%) corresponds to Sarcoptiformes. According to the taxonomic resolution, data of 170 species, 30 genera and 2 families of Collembola and 412 species, 131 genera and 49 families of Acari were obtained.

## Methods used for trophic resources identification

### Methods for resource assignment

The method of observations in laboratory tests provides the main empirical evidence from the database with 706 records (Figure 2).

**Figure 2:**
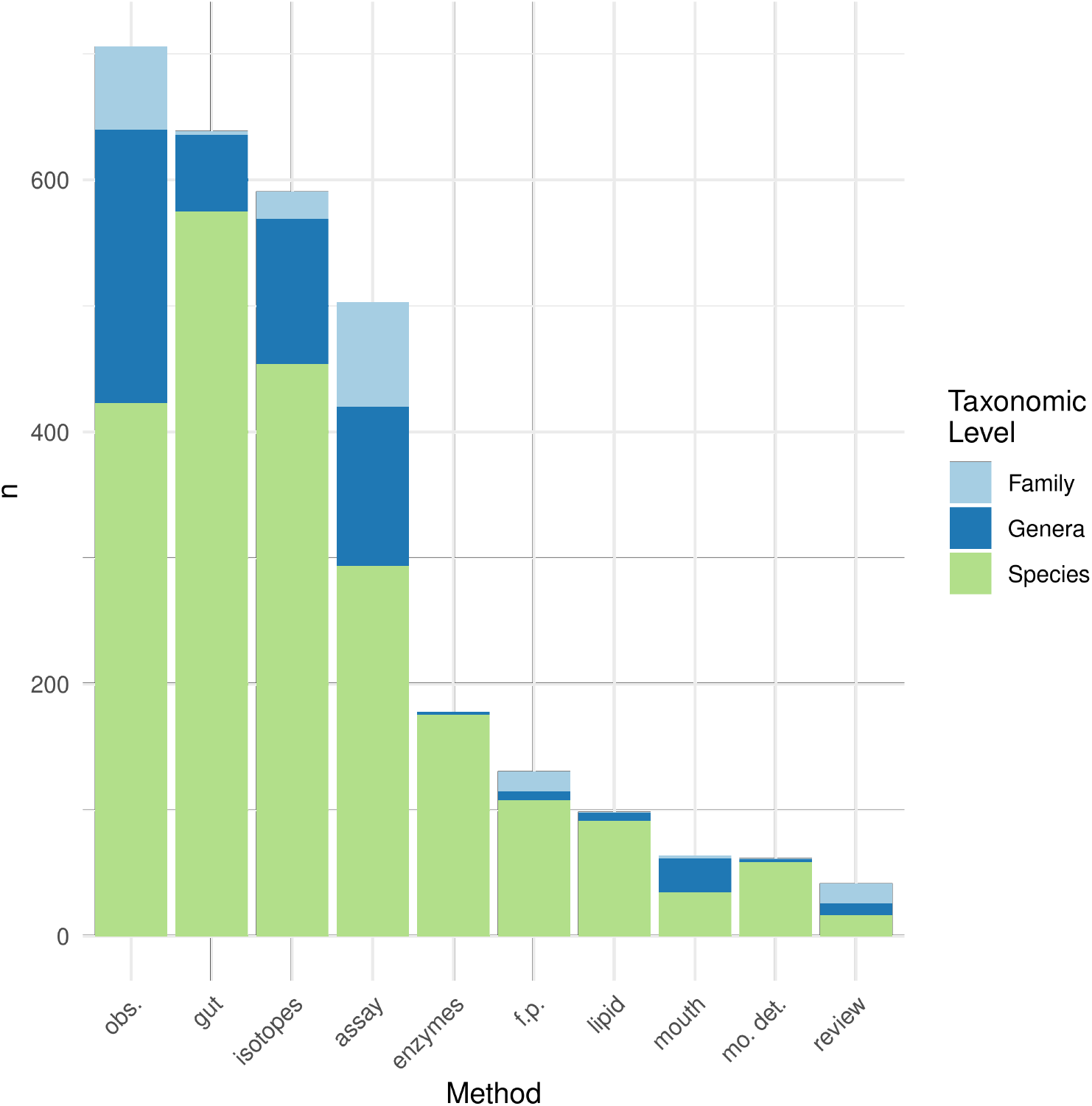
Methods used in the literature to assign trophic resources to soil mesofauna taxa. Colors within columns refer to the number of different taxonomic levels for which each method assigned at least one resource. Assay: laboratory tests and observations. Isotopes: stable isotopes. Gut: intestinal content. Enzymes: Digestive enzymes. Mouth: mouthparts morphology. Lipid: lipid profile. Mo. det: molecular detection of intestinal content, f.p.: Food preference assays. Obs.: direct lab observations of feeding activity. Reviews: general reviews by other authors.

Microarthropods at the family level constitute 9.3% of the observations in laboratory tests, with 63.6% of the total corresponding to the Prostigmata suborder, within which 10 different families are mentioned. The evidence offered by observations in laboratory tests adds up to 90.7% for species and genera. The orders Mesostigmata (62.5%), Sarcoptiformes (23.8%), Trombidiformes (9%) and Arthropleona (4%) are the ones with the higher presence in the literature.

The gut content method reaches 21.3% of the records, in which the species taxonomic level corresponds to 89.9% of the total records and the genus level to 9.5%.

Stable isotopes follow in importance. From these, 76.9% of the records mention species. Genera and family add up to 23.1% of the records.

The activity of digestive enzymes (178 records) in all cases reports down to the species level. For this empirical evidence, the authors worked with the order Sarcoptiforme (Acari) with 38 different species and Arthropleona and Symphypleona (Collembola) with 17 different species.

Other three methods accumulate 9.75% of the records, these being food preference tests (131 records), the study of fatty acids (99 records), and the use of mouthpart structures (64 records).

Intestinal contents analyses with molecular techniques for the detection of DNA is a tool of recent development and represent 2% of the total records.

### Resources identified by empirical evidence

The main resource mentioned corresponds to saprophytic fungi (19.7%) followed by microfauna (15.6%), bacteria (10.8%), and litter (10.3%), the records for mites, collembola, enchytraeidae, larvae, and eggs accumulate 725 mentions (24%) (Table 1). It is worth noting that a higher number of records have been taken to the species level. For instance, 592 trophic records are associated with saprophytic fungi consumption, of which 16 were associated with the taxonomic level of the family, 105 to the genus level, and 474 to the species level.

Laboratory observations (the most used method), mention the use of the thirteen trophic resources (Figure 3) in which the order of importance according to the number of mentions is microfauna > springtails > mites > saprophytic fungi > invertebrate eggs > enchytraeids > larvae of invertebrates that accumulate 82.3%. For this empirical method, the main resources correspond to typical resources of predatory animals except for saprophytic fungi, the main taxa mentioned is Mesostigmata. The stacked bars (Figure 3) show the different proportions in which the methods provide evidence of the use of a resource, if the contribution of each method is considered according to the number of citations in the bibliography, they are counted in decreasing order: direct observations (706 records) > intestinal content (639) > isotopes (591) > laboratory tests (503) > enzymes (178) > food preference tests (131 records).

**Figure 3:**
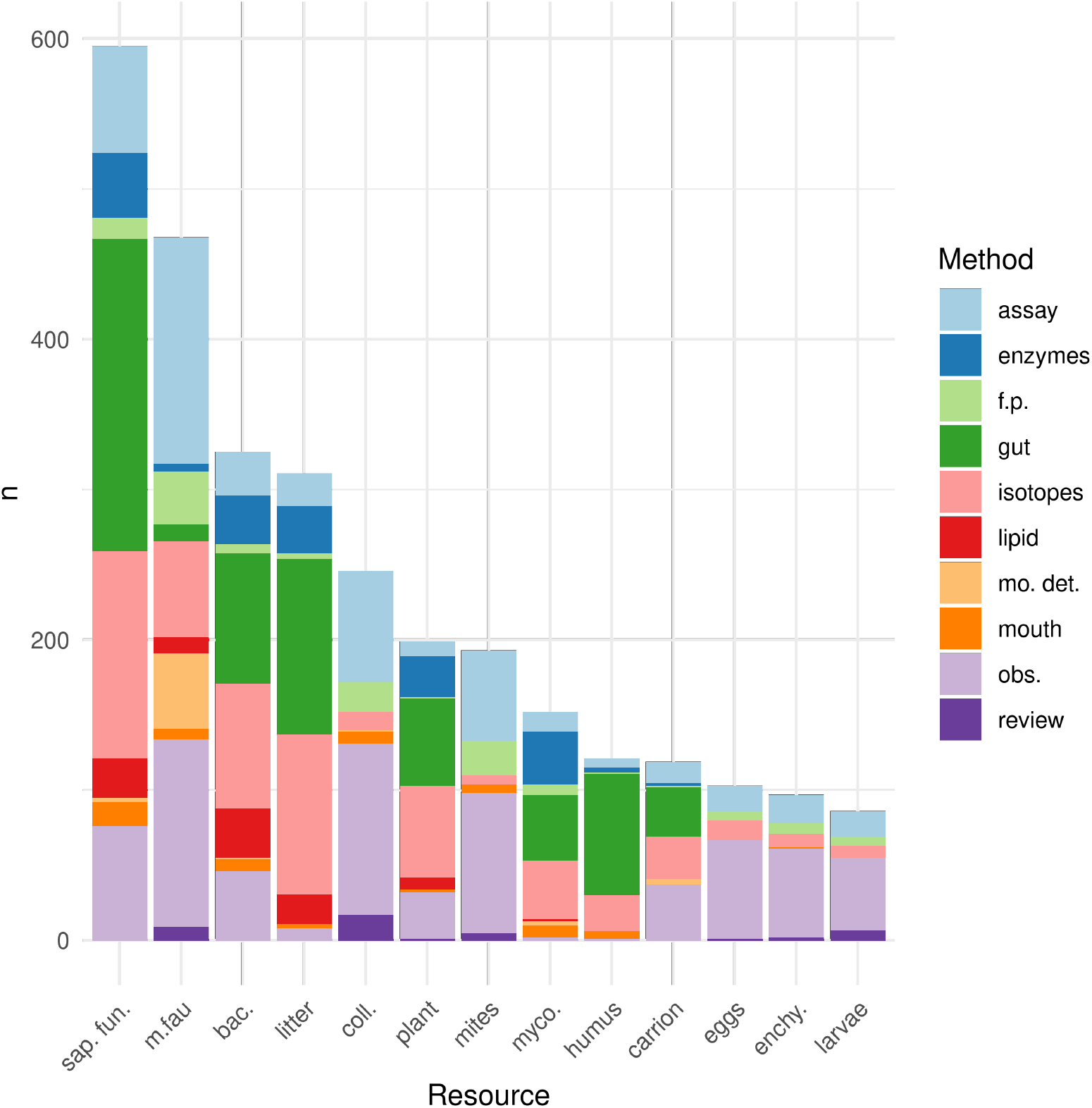
The number of records in the literature assigning each one of the 13 trophic resources to a mesofauna taxon as shown in Table 1. Colors in the columns refer to the method used to assign those trophic resources to a particular taxon. Methods as in Fig. 2.

The methodologies that use laboratory studies, i.e. laboratory tests, laboratory observations, and food preference tests, provide direct evidence of the use of the trophic resources, constituting together 44.5% of the empirical evidence analyzed. (ESM II) These laboratory methods are the ones that most frequently mention the use of animal resources, springtails - mites - invertebrate larvae - invertebrate eggs - enchytraeids, as food resources. These methods rarely mention the consumption of Mycorrhizal fungi and rarely the use of humus.

### Use of trophic resources by soil microarthropods

It is interesting to note that at the family level, the most numerous records correspond to trophic resources such as springtail, mites, microfauna, larvae, invertebrate eggs, and enchytraeids, typical resources of predatory microarthropods (Table 2). Similarly, at the genus level, the typical resources of predators represent 54% of the records. At the species level, the main resource mentioned is saprophytic fungi.

**Table 2:** Breakdown of resources by Nested boxes represent the main orders of Acari and Collembola. In parentheses is the total number of taxa for the taxonomic category. Within the cells, the number of taxa associated with the trophic resource is shown. M: Mesostigmata, S: Sarcoptiformes, T: Trombidiformes, A: Arthropleona, N: Neelipleona, Sy: Symphypleona.

| Trophic Resource | FAMILY |  |  |  |  |  | GENERA |  |  |  |  |  | SPECIES |  |  |  |  |  |
| --- | --- | --- | --- | --- | --- | --- | --- | --- | --- | --- | --- | --- | --- | --- | --- | --- | --- | --- |
|  | M | S | T | A | N | Sy | M | S | T | A | N | Sy | M | S | T | A | N | Sy |
| **Total** | 34 | 82 | 32 | 9 | 1 | 3 | 89 | 157 | 28 | 58 | 2 | 10 | 136 | 264 | 22 | 154 | 1 | 13 |
| **Saprophytic fungi** | 7 | 68 | 10 | 9 | 1 | 2 | 11 | 120 | 10 | 54 | 1 | 7 | 7 | 160 | 3 | 119 | 1 | 8 |
| **Microfauna** | 32 | 26 | 23 | 8 | 0 | 0 | 80 | 40 | 4 | 33 | 0 | 0 | 106 | 47 | 0 | 39 | 0 | 0 |
| **Bacteria** | 4 | 55 | 6 | 9 | 1 | 2 | 4 | 91 | 7 | 38 | 2 | 5 | 3 | 109 | 7 | 67 | 1 | 5 |
| **Litter** | 0 | 43 | 0 | 8 | 1 | 3 | 0 | 79 | 0 | 35 | 1 | 6 | 0 | 124 | 0 | 59 | 1 | 5 |
| **Mycorrhizal fungi** | 2 | 31 | 3 | 7 | 0 | 1 | 1 | 44 | 0 | 28 | 0 | 2 | 0 | 54 | 0 | 46 | 0 | 2 |
| **Plant tissue** | 0 | 42 | 5 | 7 | 0 | 2 | 0 | 72 | 4 | 20 | 0 | 7 | 0 | 100 | 3 | 26 | 0 | 7 |
| **Springtails** | 24 | 0 | 27 | 2 | 0 | 0 | 53 | 0 | 13 | 3 | 0 | 0 | 80 | 0 | 9 | 4 | 0 | 0 |
| **Mites** | 21 | 0 | 26 | 1 | 0 | 0 | 54 | 0 | 10 | 1 | 0 | 0 | 81 | 0 | 7 | 0 | 0 | 0 |
| **Larvae** | 12 | 0 | 9 | 0 | 0 | 0 | 32 | 0 | 0 | 0 | 0 | 0 | 41 | 0 | 0 | 0 | 0 | 0 |
| **Humus** | 0 | 24 | 2 | 8 | 1 | 2 | 0 | 27 | 0 | 26 | 1 | 3 | 0 | 28 | 0 | 49 | 1 | 3 |
| **Invertebrate eggs** | 15 | 0 | 10 | 3 | 0 | 0 | 33 | 0 | 2 | 5 | 0 | 0 | 52 | 0 | 1 | 4 | 0 | 0 |
| **Enchytraeids** | 17 | 0 | 7 | 4 | 0 | 0 | 41 | 0 | 0 | 4 | 0 | 0 | 46 | 0 | 0 | 3 | 0 | 0 |
| **Invertebrate carrion** | 5 | 22 | 2 | 6 | 1 | 2 | 9 | 30 | 0 | 28 | 1 | 3 | 11 | 31 | 0 | 40 | 1 | 4 |

The use of trophic resources in the six main orders mentioned in the literature (3 orders of Collembola and 3 orders of Acari) are divided into their families, genera, and species in a nested way (Table 2). For example, for the order Symphypleona (Sy), the empirical evidence studies 13 species included in 10 genera within 3 families. In this way, Table 2 shows also for Symphypleona (Sy), that 8 of the 13 species mentioned, within 7 of the 10 genera, within 2 of the 3 families in the available bibliography, are mentioned consuming Saprophytic fungi. For additional identity and information on families, genera and species see ESM II.

In all the taxonomic hierarchies considered, the order Sarcoptiformes (Acari) consumes mainly saprophytic fungi, bacteria, litter, plant tissues, and Mycorrhizal fungi. The order Trombidiformes (Acari), are presented mainly as predators.

For Collembola, the diversity of species addressed by empirical evidence is grouped into only 13 families, 3 of which belong to Symphypleona and a family of Neelipleona.

The empirical evidence that addresses the trophic study of Arthropleona (Collembola) is represented by 154 different species, these were mainly associated with saprophytic fungi (119 species), followed in importance by bacteria, litter and humus. The microfauna is mentioned as a resource for 39 species.

Figure 4 presents the proportion of resources used by the main 3 mite orders and 3 Collembola orders, as found in the literature. For instance, the resource of saprophytic fungi is the main constituent of the diet of the Arthropleona, being the bacteria, the litter, the humus, and the microfauna mentioned resources in lesser proportion. Trombidiformes have a diet based mainly on microarthropods and nematodes and to a lesser extent saprophytic fungi and bacteria.

**Figure 4:**
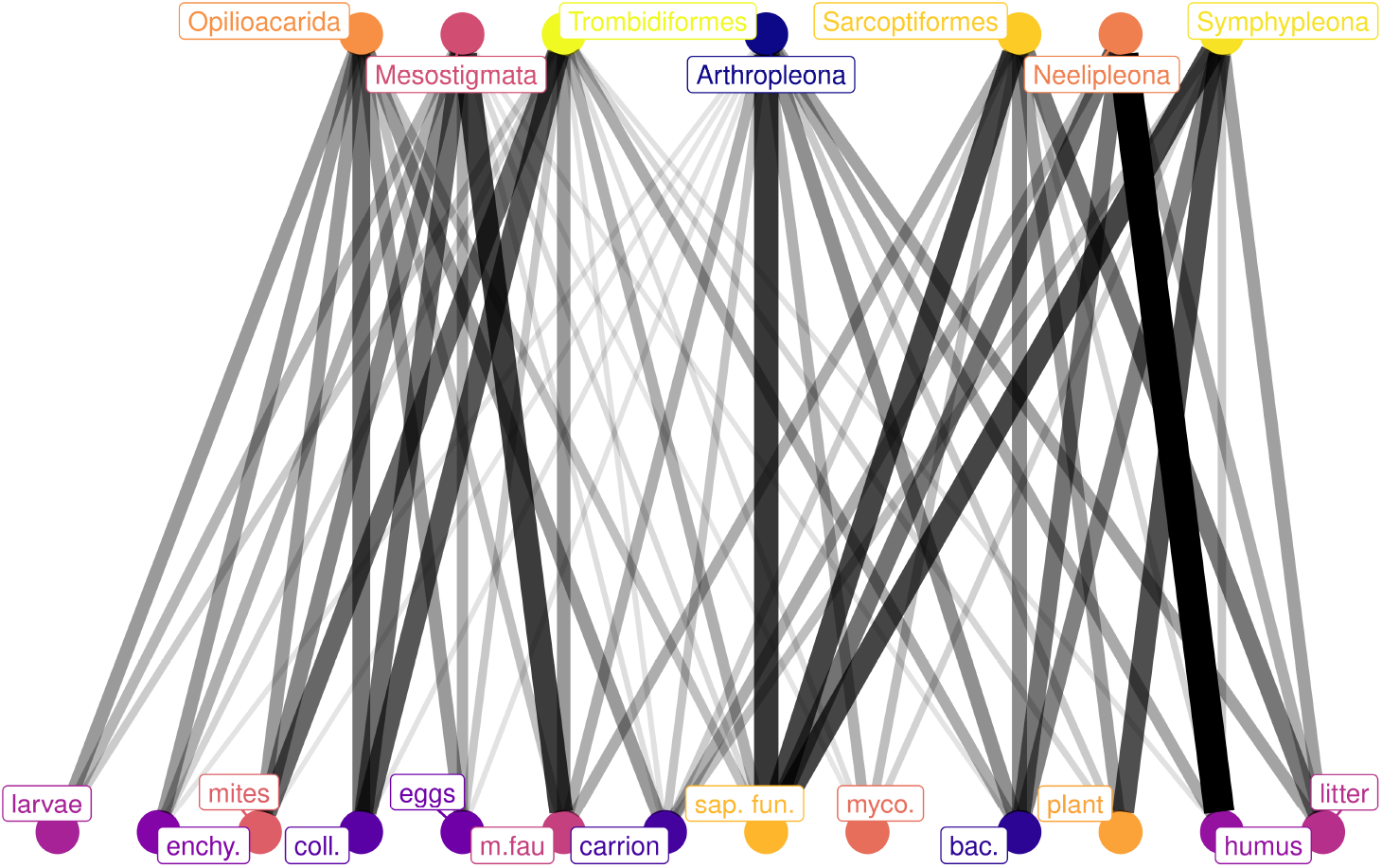
Bipartite graph showing the use of trophic resources by the main orders of Acari and Collembola. Upper nodes: Acari and Collembola orders as in Table 2. Lower nodes: trophic resources. The thickness and intensity of the lines give an idea of the proportion of mentions in the available literature about their use of trophic resources. Resource names as in Tables 1 and 2.

## Discussion

Until the 1960s, soil fauna was considered mainly earthworms, and terrestrial ecologists considered most of the soil fauna as a “black box” of decomposers and detritivores (Briones, 2014). The information gradually collected over the following decades regarding the edaphic microarthropods is reaching the point where it is possible to focus on integrative works. (Pautasso 2013) However, the available information is still scarce and mostly restricted to the most numerous or conspicuous groups of the soil microarthropods, mainly from European soils.

For soil microarthropods, the evidence provided by laboratory work results in the most straightforward traditional method to know how these animals use trophic resources, their feeding behavior, their food preferences, and their development and growth in controlled environments, but Feeding links observed in the laboratory do not necessarily relevant for the field (Nielsen 2018, Potapov 2022). The method used to relate the isotopic signature of organisms and their resources is a recently developed tool that is useful for detecting the importance and the changes in time or space of assumed trophic relationships, which means that we have to know in advance if the trophic relationship exists. The main drawback of the method is that if we erroneously assign a trophic relationship the proportions of the diets could be greatly distorted from the real trophic relationships. On the other side, this method has several advantages 1) it can analyze a large number of species and an important variety of resources (Potapov et al., 2019), 2) it can provide field evidence of real interactions

3) it can provide quantitative data, accounting for interaction strength
4) it can provide information on assimilation and not only ingestion (Nielsen et al. 2018)

The different methods have different sources of errors so it will be desirable that trophic resources used by soil microarthropods can be determined in a complementary way with several methods (Potapov et al., 2020; Walter et al.,1991).

The most used resources are saprophytic fungi, microfauna, and bacteria. If we associate them with their nutritional characteristics, these trophic resources are rich in molecules with great nutritional value as determined in the dietary routes labeled by fatty acids, stable isotopes, or the enzymatic methods (Nielsen, 1962; Potapov et al., 2019; Ruess and Chamberlain, 2010).

It is also to be noted that the information available for Acari and Collembola is strongly asymmetric, corresponding mainly to the order Sarcoptiformes in Acari and to Arthropleona in Collembola.

Despite the increasing amount of descriptive works and lists of taxonomic groups, the information available worldwide is still largely fragmentary and incomplete, and taxonomic resolution varies considerably between and within published works.

We found that a large proportion of the resources are defined as taxonomic categories of species, genera, and families, which would be important to estimate the diets of higher taxonomic groups. The available information for low taxonomic levels could be used as a reference to address the problem of what the use of soil resources will be like by higher-level taxonomic groups (Bedano, 2007; Potapov et al. 2020).

However, this information must be interpreted with caution, because within a taxonomic category each species could apply different strategies when exploiting food resources (Lavelle and Spain, 2001; Moore et al., 1988), although the taxa treated here are considered generalist consumers, recently the term “choosy generalist” was suggested as the behavior that characterizes consumers that inhabit soils (Potapov et al. 2022)

The results presented here provide valuable new information about the different feeding strategies of the main groups of the soil microarthropods, and also on the quality and usefulness of the different methods used to assign trophic resource use to different taxa. It also presents the current status of knowledge about soil microarthropods trophic resources usage. Moreover, it highlights the still quite scant information available in this regard.

Finally, it is necessary to call attention to the need for more studies on the trophic relationships of the soil microarthropods. Out of a total of approximately 9000 described species of Collembola (Bellinger et al., 2020), it was only possible to find trophic information for just 127 soil inhabitant species. For Acari, out of the approximately 58,000 species described (Schmidt, 2020), it was only possible to find references of trophic relationships for 307 soil species.

It is clear that big gaps in the available information must be filled to advance our knowledge on the structure and functioning of soil food webs. Not only are there several microarthropods groups hardly explored, but also the geographical coverage is still quite narrow. Almost 52% of the published studies about the trophic resources of Acari and Collembola were developed in European countries. Further deepening on the knowledge of functional and trophic relationships of the soil the fauna would allow for a better and more precise evaluation of the functioning of the edaphic ecosystem, the protection of the ecosystem services that the soil microarthropods provide, and the sustainable use of the soil.

## Supporting information

Supplementary Material III

## Acknowledgements

The authors wish to thank the help of Dr. Fernando Momo for his useful comments to previous versions of this work. To Dr. Mark Breidenbaugh for his help in reviewing the English language usage and grammar.

## Declarations

This work was partially funded by a scholarship to Victor N. Velazco from Consejo Nacional de Ciencia y Tecnología (CONICET). This research did not receive any other specific grant from funding agencies in the public, commercial, or not-for-profit sectors. The authors declare no conflicts of interest. All the data for this work not shown here are available as Electronic Supplementary Material and referred to in the text when appropriate.

## [Supplementary Material Captions]

Electronic Supplementary Material I (ESM I): Complete list of bibliographic sources that relate specific taxa to a particular trophic resource.

Electronic Supplementary Material II (ESM II): Complete list of 2717 taxa for which a particular trophic resource has been assigned.

Electronic Supplementary Material III (ESM III): Flowchart showing the criteria used in the bibliographic search and selection.

