## Supplementary Material III for "Trophic resources of the edaphic microarthropods: a worldwide review of the empirical evidence"

### Supplementary material III: Bibliography search and paper selection process flowchart

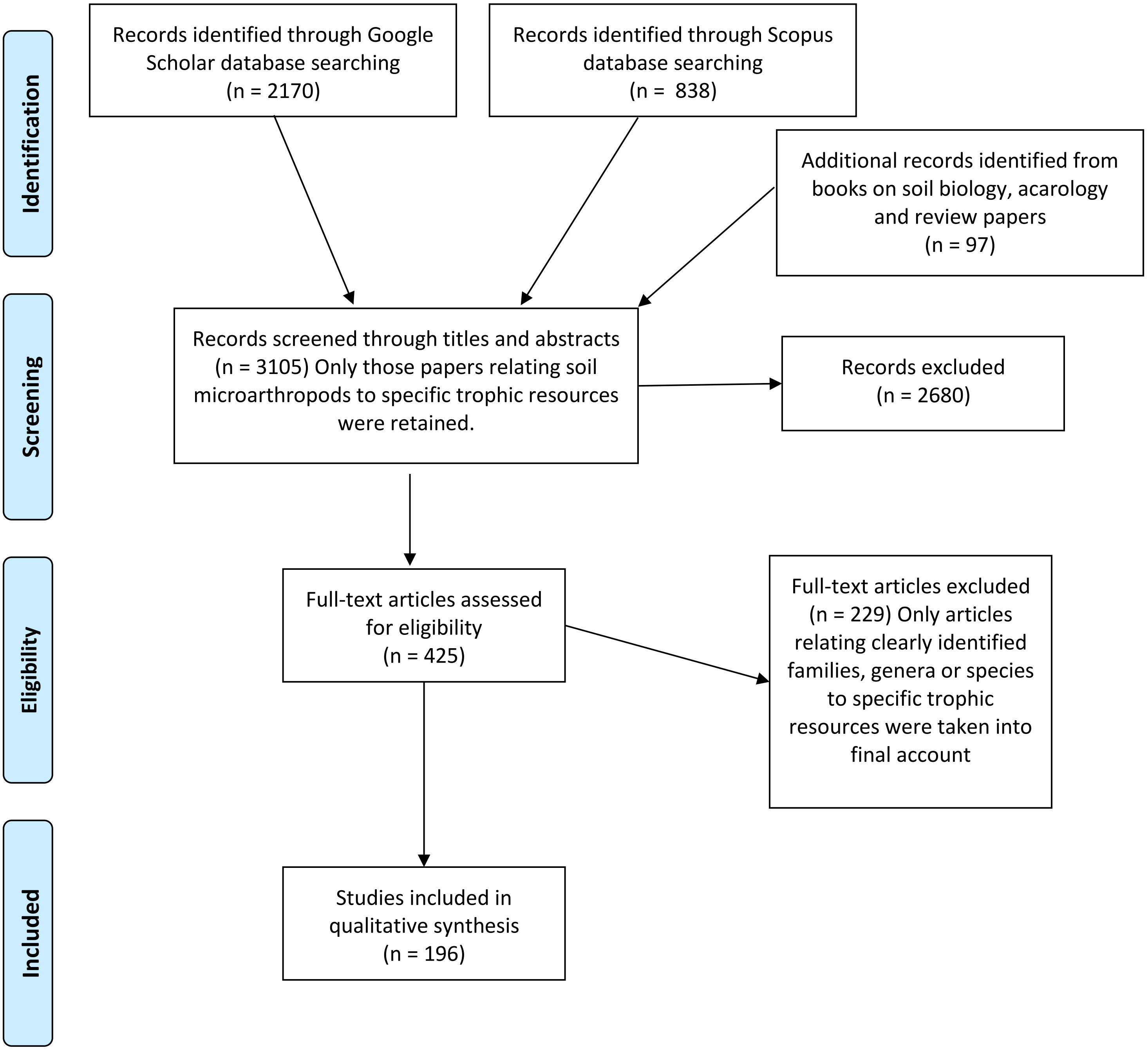


*From:* Moher D, Liberati A, Tetzlaff J, Altman DG, The PRISMA Group (2009). *P*referred *R*eporting *I*tems for *S*ystematic Reviews and *M*eta-*A*nalyses: The PRISMA Statement. PLoS Med 6(7): e1000097. doi:10.1371/journal.pmed1000097

**For more information, visit www.prisma-statement.org.**
